# GEGVIC: A workflow to analyze Gene Expression, Genetic Variations and Immune cell Composition of tumor samples using Next Generation Sequencing data

**DOI:** 10.1101/2023.04.05.535678

**Authors:** Oriol Arqués, Laia Bassaganyas

## Abstract

**Background:** The application of next-generation sequencing techniques for genome and transcriptome profiling is to build the main source of data for cancer research. Hundreds of bioinformatic pipelines have been developed to handle the data generated by these technologies, but their use often requires specialized expertise in data wrangling and analysis that limit many biomedical researchers. Providing easy-to-use, yet comprehensive and integrative open-source tools is essential to help wet-lab and clinical scientists feel more autonomous in performing common omics data analysis in cancer research.

**Results:** Here, we present GEGVIC, an R tool to easily perform a set of frequently used analyses in cancer research, including differential gene expression, genomic mutations exploration and immune cell deconvolution using minimally processed human/mouse genomic and transcriptomic sequencing data. GEGVIC is designed as a modular pipeline that combines a variety of widely used available methods distributed in three principal modules (*Gene Expression*, *Genomic Variation* and *Immune Composition*), which run independently and include several visualization tools. This open-source software is also presented as a graphical user interface (GUI) using the Shiny framework (*GEGVICShine*) to eliminate the coding barrier for non-R users and enable comprehensive analyses of tumor samples via one-click features.

**Conclusions:** In summary, GEGVIC provides a simple, powerful and highly flexible workflow for researchers to process and interpret tumor transcriptomic and genomic data while decreasing or eliminating coding burden and facilitating efficiency for inexperienced bioinformatics users. GEGVIC R package instructions and source code are published on Github (https://github.com/oriolarques/GEGVIC), whereas GEGVICShine is hosted at https://gegvic.vhio.net/.

## Background

Research in oncology increasingly relies on high-throughput technologies to dissect the complex biology of tumors and the tumor microenvironment (TME). DNA sequencing (DNA-seq) and RNA sequencing (RNA-seq) have become central tools in the cancer field, where they are used to uncover the molecular mechanisms underlying the disease, as well as the dynamics of tumors and TME based on their genomic and transcriptomic profiles. Indeed, the application of such omics technologies to primary and metastatic tumor analyses has already resulted in notable advances in our fundamental understanding of the biological landscapes of most tumor types, with numerous discoveries relevant to cancer diagnosis and treatment (1,2).

The steady progresses of sequencing technologies and the rapid accumulation of massive amounts of omics data have catalyzed the development of a plethora of bioinformatics tools for the analysis of complex, high-dimensional datasets that are continuously being developed and improved (3,4). This poses considerable challenges for data processing and analysis, and often require additional steps to achieve meaningful results. In addition, most algorithms are limited to experienced bioinformatics users and require significant computational resources. While this does not pose a problem for bioinformatics teams and research groups located in large collaborative academic environments, for the large majority of experimental laboratories, the day-to-day bioinformatic skills and computational infrastructure available are limited to a small number of analyses. This creates a significant operational disequilibrium, as laboratories with direct access to data from pre-clinical animal models and/or patient samples with valuable clinical information often lack the ability to analyze them. A common solution is to establish collaborations with core facilities or industry partners not only to generate the raw data but also to conduct analyses, which comes with some limitations. On the one hand, not all studies, types of data, and contexts fit into the standard and fixed pipelines normally used in these collaborations, and downstream analyses often requires several iterations that could be more efficiently performed by the researcher who designed the experiment. On the other hand, in situations such as preliminary analyses, public data exploration or hypothesis generation, these collaborations are not cost-effective. The alternative of commercial bioinformatics solutions exists, but the cost of software access may not be affordable for many academic laboratories. As sequencing technology is now undeniably a potent driver of scientific development even in molecular or clinical setting, providing effective user-friendly open-source tools that can help to bridge the gap between the investigator’s abilities to generate and analyze data is therefore essential.

Omics data inform on many dimensions of cellular systems and biological processes which can be differently inferred from genomic or transcriptomic data. This justifies the wide range of bioinformatic techniques being developed to answer complex cancer-related questions, ranging from how tumors evolve to how to better predict clinical outcomes (3). However, the most common starting-point for cancer studies from high-throughput datasets is based on defining a set of different molecular features for phenotypes of interest from bulk genomic and/or transcriptomic readouts. This includes the quantification and description of the somatic mutational landscape, the identification of differentially expressed genes between groups (Differential Gene Expression or DGE analysis) along with the signaling pathways they target, and the assessment of the immune cell composition and density in the TME. Methods are already available to easily perform some of these analyses (5–11). For instance, Shiny-Seq (10) and GENAVi (11) are both shiny web applications specifically developed for comprehensive gene expression analyses including DGE and pathway enrichment; MutationExplorer (7) uses bulk RNA-seq data to explore cancer somatic mutation landscapes; MuSiCa (8) and signeR (9), also available as a shiny app (signeRFlow), are user-friendly methods for mutational signature discovery from genomic data; and the recent MouSR (5) enables users to easily interrogate gene expression data from bulk RNA-seq to perform DGE, gene set enrichment, and TME cell population analyses from human or mouse tissue. However, all of them are independent applications and the integration of their results is not straightforward.

In order to simplify and improve the accessibility of such multidimensional (genomic and transcriptomic) bioinformatics analyses, we created a modularized R package named GEGVIC (Gene Expression, Genetic Variations and Immune cell Composition). This package integrates the most frequently used approaches to analyze human or mouse transcriptomic and genomic data in cancer research. GEGVIC can be executed by functions as an R package or as a graphical user interface (GUI) using the Shiny framework (*GEGVICShine*). GEGVIC has been developed with the aim of facilitating the execution of comprehensive analysis of tumor samples and their microenvironment’s immune composition. By eliminating or decreasing the coding burden for inexperienced bioinformatics users, our package allows them to focus on the biological questions and data interpretation. We improve on available tools by offering integrated, simply accessible and easily expandable tools for the most common set of cancer-related analyses, which reduces the need to use multiple independent analytical packages in order to obtain a complete picture of the basic aspects required for any cancer study.

## Implementation and applicability

GEGVIC comprises three main modules that run independently: 1) *Gene Expression,* 2) *Immune Composition* and 3) *Genomic Variation* (**Figure 1**). Modules 1 and 2 use transcriptomic data whereas module 3 uses genomic data. Inputs consist of minimally processed transcriptomic (read counts table) or genomic mutations (*maf* format table) data, which are the universal format files produced by most alignment and variant calling pipelines. A metadata file containing one or more categorical variable of interest for each sample (e.g., treatment or response status, absence/presence of a genomic alteration, high/low aneuploidy burden) is also required to define groups or conditions. As an R package, the tool can be executed function-by-function or, alternatively, module-by-module, by using a single module-specific function. Plots can be customized by modifying specific parameters incorporated within the functions. As a web-based tool, users may select their modules of interest, after which they are intuitively guided through the different steps necessary to perform the analyses and customize the data plots. Results from the web-interface are the same as the ones obtained from the R-package, with the additional advantage that the tables generated with the GUI are interactive.

**Figure 1.**
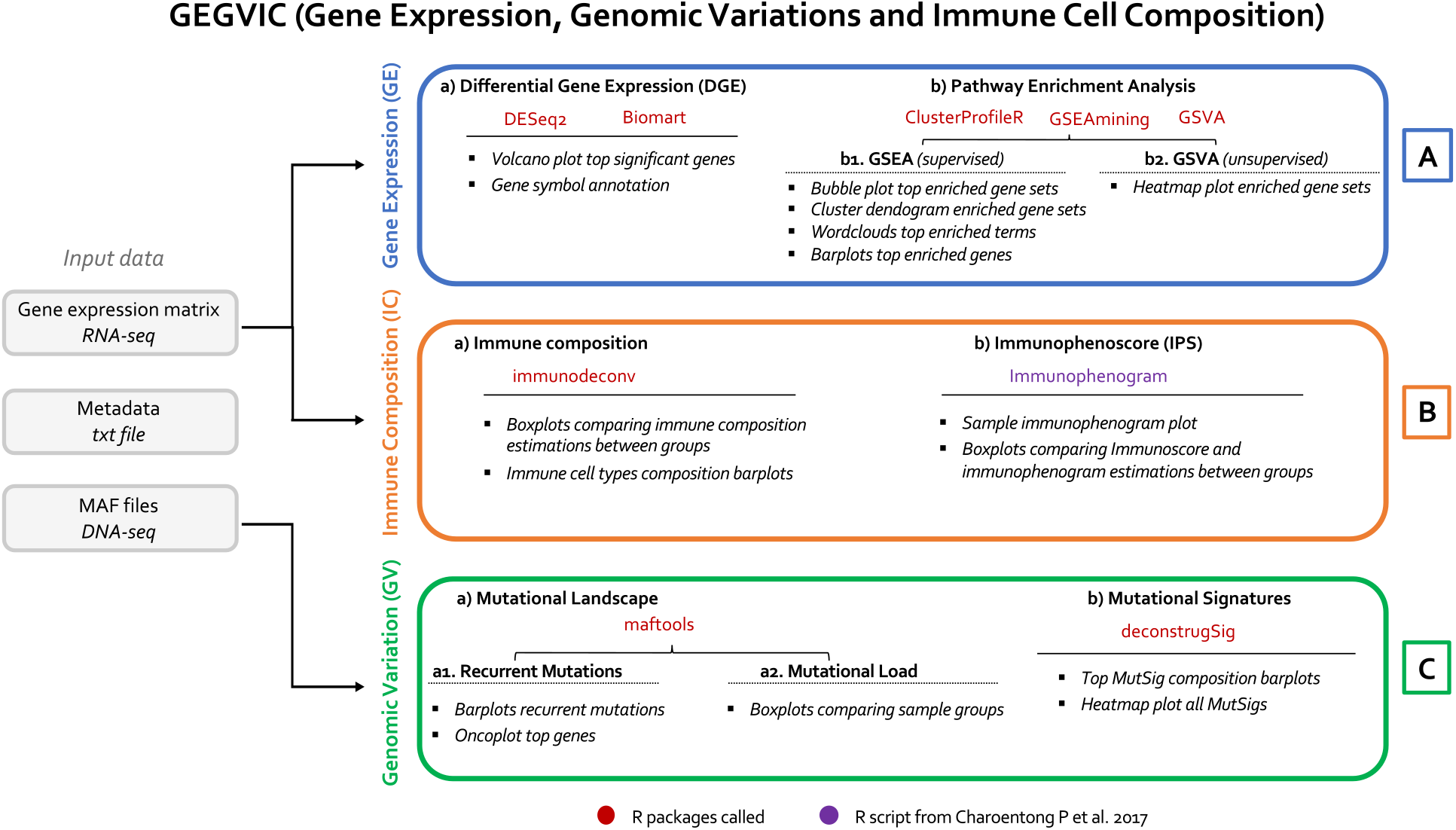
GEGVIC modular framework. GEGVIC is divided in three main analytical modules that n independently and include commonly used R packages (red and purple). Gene expression an be used for *Gene Expression (GE)* and *Immune Composition (IC)* modules, while ic data (MAF file) is exclusively for the *Genomic Variation (GV)* module. The metadata file is d for all modules. Proposed approaches are highlighted in bold and the outputs generated hlighted in italics.

The software is also adapted to the analysis of mouse (*Mus musculus*) data. Although some methods integrated in our tool were designed to work solely with the human genome, GEGVIC automatically converts mouse gene symbols to their human orthologues through the functionalities of the BioMartR package (12). A comprehensive description of the tool is provided herein, where each module is detailed specifying the utility of each function from the R package and the outputs generated.

## 1. Gene Expression module (GE)

This module uses the functionalities provided by DESeq2 (13), with the default parameters for data normalization recommended for bulk RNA-seq. The two main analyses included are (a) the computation of the Differential Gene Expression (DGE) between groups of interest and (b) the subsequent pathway enrichment analysis of genes found by the DGE (**Figure 1A, Supplementary Figure 1**). Before the computation of the DGE, GEGVIC facilitates a first data exploration by performing a principal component analysis (PCA) using the gene expression counts through the function `ge_pca()`. This function follows DESeq2 steps to normalize counts and transform them using the variance stabilizing transformation (VST).

The DGE analysis comparing the selected groups of interest is computed using the `ge_diff_exp()` function. The function allows the user to select which logFC shrinkage method to apply (see *Supplementary Material*). Multiple comparisons are possible when variables contain more than two levels. The output is saved in an object (named *results.dds*), which can be consulted by accessing the results table and/or visualized using the `ge_volcano()` function. By default, the ten most significantly up- and down-regulated genes are highlighted. If needed, the ENSEMBL or NCBI gene identifiers can be directly converted to HGNC symbols by running the `ge_annot()` on the *results.dds* object and by specifying the Biomart database of interest. The reference genomes from the two last versions of human and mouse genomes (GRCh37 and GRCh38; GRCm38 and GRCm39, respectively) are built in the tool, but the user may also incorporate other versions by previously downloading them.

Next, to perform the functional enrichment analysis from the genes identified by the DGE, GEGVIC provides two complementary methods: Gene Set Enrichment Analysis (GSEA) (14) and Gene Set Variation Analysis (GSVA) (15). The GSEA uses permutation approaches to establish whether a specific pre-defined gene set is significantly enriched in one of the comparing groups of interest (*supervised* comparison analysis). In contrast, the Gene Set Variation Analysis (GSVA) estimates the variation of pathways’ activity over a dataset by translating, for each sample, the gene expression information to gene set enrichment scores, which allows to evaluate the degree and/or type of clustering between samples in an *unsupervised* manner. Both methods require a collection of gene sets provided by the user, which must be encoded within a single file following the GMT (Gene Matrix Transposed) format. We recommend using collections from the Molecular Signatures Database (MSigDB), one of the most comprehensive databases of gene sets for performing functional enrichment analysis (16). For further information see *Supplementary Material*.

To perform the GSEA, the `ge_gsea()` function takes the functionalities of clusterProfileR (17), with the option of modifying the adjusted p-value cutoff if needed. The top regulated gene sets (20 up- and 20 down-) according to the Normalized Enrichment Score (NES) are represented as a bubble plot. To facilitate the interpretation of all the significant information obtained, the same gene sets are grouped by similarity and plotted using the GSEAmining R package (18). These plots include: (i) a dendrogram clustering the gene sets; (ii) a word-cloud of biological terms enriched at each cluster; and (iii) the top three genes in the leading-edge analysis included within the enriched gene sets present in the cluster. To perform the GSVA, the `ge_single()` function uses the GSVA R package (15). The GSVA results are presented in a form of a clustered heatmap highlighting the groups of interest and the enrichment score of each gene set. This unsupervised clustering of samples using biological pathways instead of genes is complementary to the PCA and can be used to determine the biological relationship between the samples and groups. In both cases, output table can be saved in specific objects (gsea.res or gsva.res) and consulted or exported afterwards.

## 2. Immune cell Composition module (IC)

This module allows to infer the cellular composition and status of immune infiltrates in the TME for each sample and to compare the results obtained for specific groups of interest (**Figure 1B, Supplementary Figure 2**). Since TME cell deconvolution from bulk transcriptomes is technically challenging and the current algorithms vary in their advantages and pitfalls (19), GEGVIC includes different methodological approximations that generate a wide range of comparable and/or complementary results at the same time, giving a more complete and confident picture to the user. Note that before running these analyses, the RNA-seq count data must be transformed to transcripts per million (TPM) using the `ic_raw_to_tpm()` function.

First, an estimation of immune cell fractions for each sample can be obtained by performing the comprehensive analysis provided by the immunedeconv R package (20) and incorporated in our tool through the `ic_deconv()` function. This package includes six commonly used computational methods (quanTIseq (21), TIMER (22), MCP-Counter (23), xCell (24), EPIC (25) and CIBERSORT (26)) that are run automatically at once except for the CIBERSORT, which requires a previous download and storage of the source code in a user-defined path that needs to be specified (see *Supplementary Material* or GitHub guide for further information). Results are saved in the *ic.pred* object, which is a table containing the raw score/abundances assigned to each cell type by each method for each sample.

These approaches differ in their scoring strategies as well as the number and names of immune cell populations they examine, making direct comparison and interpretation of results difficult. For instance, xCell predicts 39 immune cell types including a wide range of very specific subtypes, whereas TIMER only predicts 6 general cell types. Although all results are available through the *ic.pred* object, GEGVIC also incorporates the `ic_plot_comp_samples()` function, which allows the integration and visualization of the data to facilitate a first global interpretation of the immune composition of the analyzed datasets. The function groups the immune cell subtypes in seven major categories (B-Cells, Macrophages, mDendritic cells, Neutrophils, NK-Cells, T-CD4+ and T- CD8+ cells), and plots the distribution of the scores obtained for each comparison group and method for each immune category (*Supplementary Material*). The statistical method for comparison of means (‘t-test’, ‘wilcox.test’, ‘anova’ or ‘kruskal.test’) can be defined by the user according to the type of data and number of groups to be compared. Besides this, given that CIBERSORT, EPIC and quanTIseq also calculate the absolute cell fractions of each immune cell population within each sample, our tool enables to plot these results through the `ic_plot_comp_celltypes()` function.

Finally, to obtain a complementary approximation of the functional state of the immune infiltrates, GEGVIC incorporates the ‘Immunophenoscore’ (IPS) approach developed by Charoentong et al. (27) through the `ic_score()` function, which uses a defined set of genes to calculate an objective score reflecting the global functional state of the immune system in each sample. The function plots the distribution of the scores obtained for each comparison group and the statistical difference can be inferred, as before, with the method selected by the user. Additionally, to enable a more comprehensive picture of the active immune infiltrates types, distribution plots are also generated for the four major determinants of the immune response used to calculate the overall IPS, i.e. MHC molecules (MHC), Checkpoints-Immunomodulators (CP), Effector Cells (EC) and Suppressor Cells (SC), and an ImmunoPhenoGram (IPG) displaying the average expression of those genes defining the different immune subcategories is provided for each sample in a pdf output file.

## 3. Genomic Variations module (GV)

Unlike the two previous modules, this one uses DNA-seq data in order to complement the transcriptomic results with the assessment of specific genomic mutations (**Figure 1C, Supplementary Figure 3**). Taking the standard *maf* files obtained from any DNA variant caller as input, the two main analyses comprised here consist of: (a) an overview of the mutational landscape identified in the studied cohort and comparison groups by using the functionalities of maftools (28); and (b) a mutational signature analysis of each sample provided by the deconstructSig package (29).

In terms of mutational landscape, two key aspects include: (i) the characterization of those alterations found at higher frequency than expected in a given dataset or comparison group, which are generally interpreted as ‘driver’ events essential for neoplastic transformation (30); and (ii) the impact of accumulating mutations in the tumor genome (*Tumor Mutation Burden* or TMB), which may be indicative of genomic instability and has also been linked to immunotherapy response (31,32). To facilitate the analysis of these aspects, GEGVIC incorporates two functions. The first one consists of the `gv_mut_summary()`, which calculates the number of genomic variants present in each sample and the frequency of each mutation in the study cohort. To display the results, the function generates an oncoplot containing a given number of the top recurrently mutated genes (defined by the user), accompanied by several barplots providing general information on the type of variants detected in the dataset. Within the oncoplot, the samples are sorted by the established comparison groups, facilitating the comparative assessment of their mutational landscape. The second one, the `gv_mut_load()` function, plots the distribution of mutational load (i.e., the total number of mutations in each sample) for each group of interest and performs the statistical comparison of these distributions using a user-defined method.

The additional genomic approximation consists in predicting the mutational signatures contributing to each tumor sample (33). To do so, the `gv_mut_signatures()` function calculates the weight of each mutational signature in each sample and generates two complementary outputs: first, a typical proportion barplot split comparing groups showing the four signatures that contribute the most to the mutational pattern of each sample; and second, a heatmap, also split by groups of interest, in which the intensity of the color is determined by the weight of each signature in each sample. Although the first one is more commonly found in the literature, the latter may appear clearer when several different mutational signatures are present in a large number of samples. Note that the function requires the user to define the version of the human or mouse reference genome to work with (i.e., GRCh37, GRCh38, GRCm38 or GRCm39), as well as the COSMIC mutational signature matrix of choice (v2 or v3.2). See https://github.com/oriolarques/GEGVIC for further information.

## Results

To exemplify the applicability of GEGVIC in a biologically relevant scenario, we used the publicly available genomic and transcriptomic data from the TCGA colorectal cancer dataset (COAD- READ), and we established the annotated microsatellite instability (MSI) / microsatellite stability (MSS) tumor status as comparative variables. A random subset of this data is available as a *demo* input within the package.

MSI tumors represent a well-defined subset of colorectal carcinomas (CRC) with special molecular etiology and characteristic clinical features, including an increased survival rate and higher sensitivity to immune checkpoint blockade (34,35). They are characterized by a hypermutated phenotype caused by incompetent DNA mismatch-repair (MMR), and typically present high levels of immune cell infiltration (35–37). Consistently, the mutational analysis from the *Genomic Variation* module showed (a) a significantly higher mutational load of MSI tumors compared to the MSS tumors (**Figure 2a**); (b) the oncoplot illustrating the genetic alterations typically associated with CRC (e.g. *APC, TP53, SMAD4* and *PIK3CA*), where is it also possible to visualize the expected lower proportion of *APC* mutations and the higher proportion of *BRAF^V600E^* mutations in MSI-related tumors, as well as their enrichment in *TTN* mutations, a pan-cancer biomarker of high tumor mutation burden (38) (**Figure 2b**); and (c) the enrichment of signatures 6 and 15, associated with deficiencies of MMR (39), in the MSI tumors (**Figure 2c** and **2d**).

**Figure 2.**
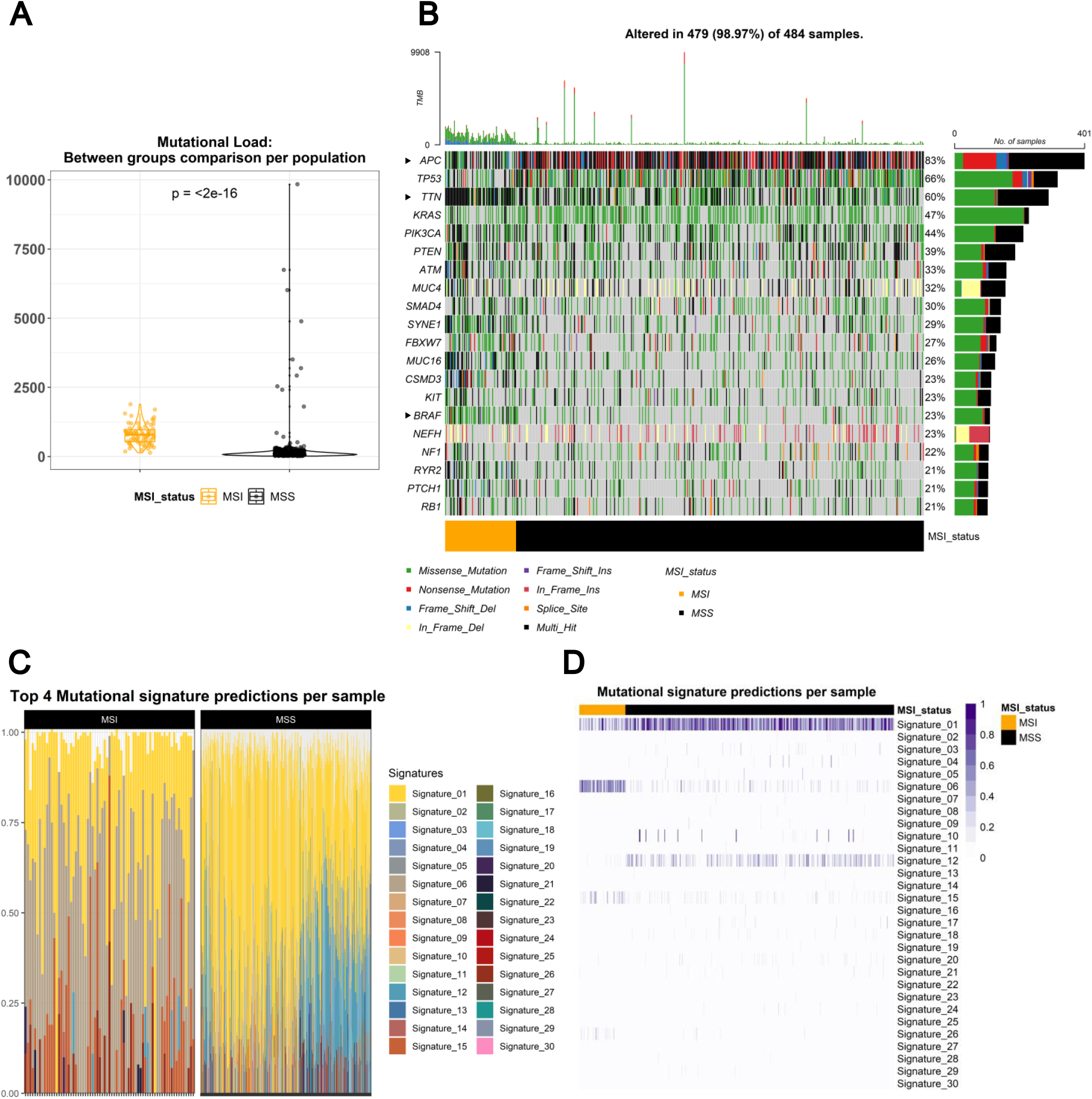
Mutational landscape of MSI and MSS colorectal tumors. Mutational analysis from ic Variation (GV) module generated (**a**) a boxplot showing the significant higher mutational MSI tumors in comparison to MSS; (**b**) an oncoplot displaying the most common mutations ed in the cohort, ordered by the MSI / MSS variable; (**c**) a stacked barplot visualizing the ion of the top 4 mutational signatures (COSMIC v2) identified in each subset; and (**d**) a p showing the enrichment of each mutational signature in each sample, ordered by the MSI variable.

In terms of transcriptomic data, the expression analysis from the *Gene Expression* module provides (a) the PCA obtained from the gene-count matrix which discriminates the MSI and MSS groups (**Figure 3a**), directly indicating the global biological distinction between both tumor subtypes; (b) the volcano plot highlighting the top-10 significantly upregulated and the top-10 signficantly downregulated genes in MSI in comparison with MSS (**Figure 3b**); and (c) the expected enrichment of terms and pathways related to immune system modulation in MSI tumors, observed from both GSEA and GSVA analyses, as well as the downregulation of processes related to lipid metabolism that is also in consistence with previous findings (40) (**Figure 3c** and **3d**). Finally, the *Immune Composition* module allowed to better explore to which extent the immune landscape of the TME differs between MSI and MSS tumors (**Figure 4**). Interestingly, CRC patients with an MSI phenotype typically display a high levels of T-lymphocyte infiltration and strong antitumor immune response, which has been associated with the high mutational burden and neoantigen load (41,42). Instead, MSS CRC harbors a disrupted immune response in the TME, characterized by low infiltrated T cells and reduced immune cytotoxicity (43). Consistent with these observations, the application of the GEGVIC integrative strategy to the *immunodeconv* results revealed a significant enrichment in CD4+T / CD8+T lymphocytes and macrophages in the MSI tumors (**Figure 4a**) (44,45). Similarly, the results from the IPS analysis indicated that the higher immunogenicity of the MSI samples can be explained by the higher proportion of immune effector cells (i.e., CD4+ and CD8+ T-lymphocytes) and the stronger expression of MHC-related molecules (**Figure 4c**). In contrast, the MSS samples showed an enrichment of immunosuppressive cells, represented by regulatory T (Treg) lymphocytes and myeloid-derived suppressor cells (MDSCs) (see *Supplementary Material*).

**Figure 3.**
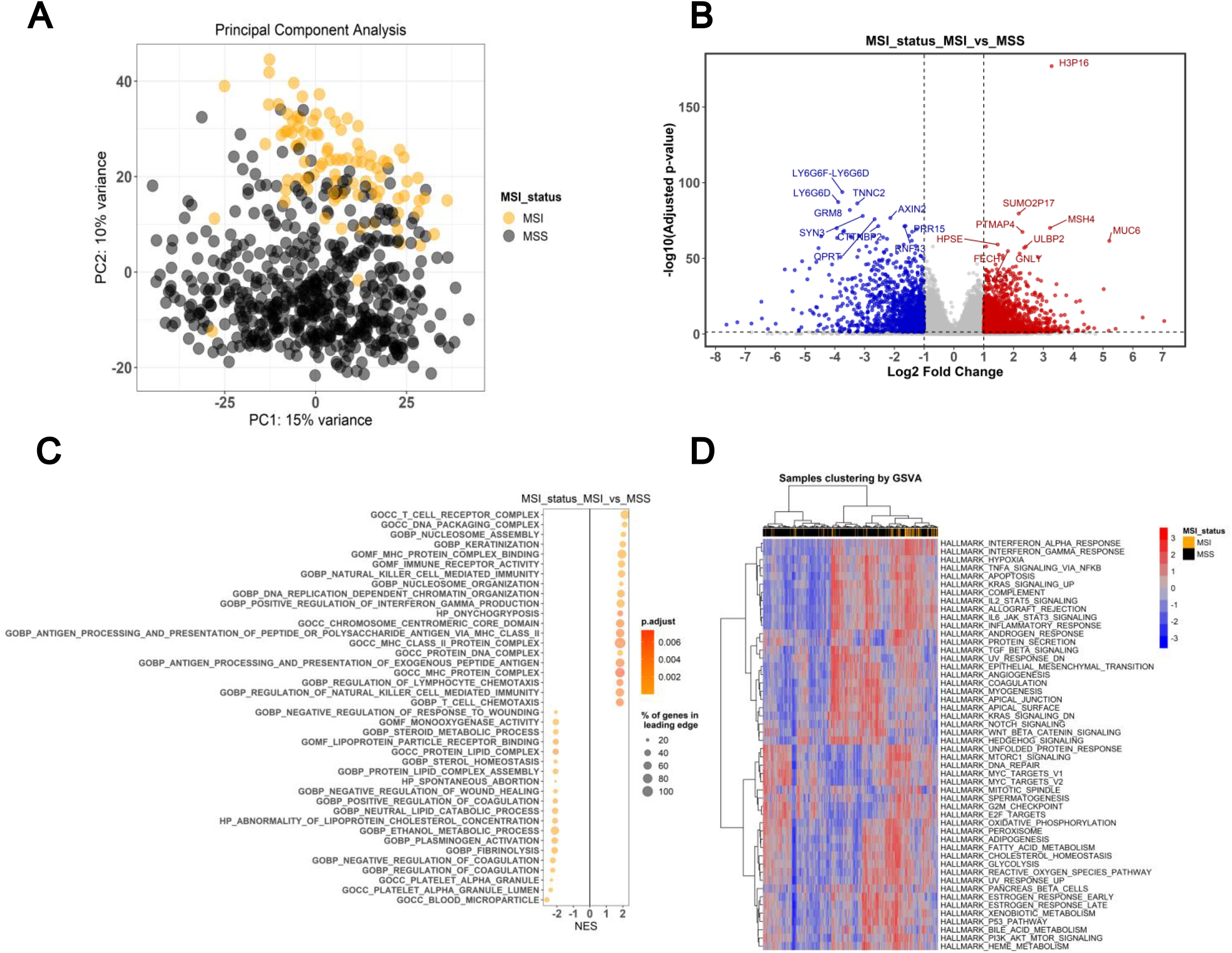
Differential gene expression and pathway enrichment analysis in MSI and MSS ctal tumors. Gene Expression analysis from Gene Expression (GE) module generated (**a**) a ot highlighting the MSI / MSS variable; (**b**) a volcano plot showing the top 10 up- and top 10 egulated genes obtained from the differential gene expression analysis between MSI and MSS; **c**) a bubble plot with the top 20 enriched GO pathways in the comparison between MSI / MSS; and (**d**) the heatmap of normalized GSVA enrichment scores for the HALLMARK gene sets on for each sample.

**Figure 4.**
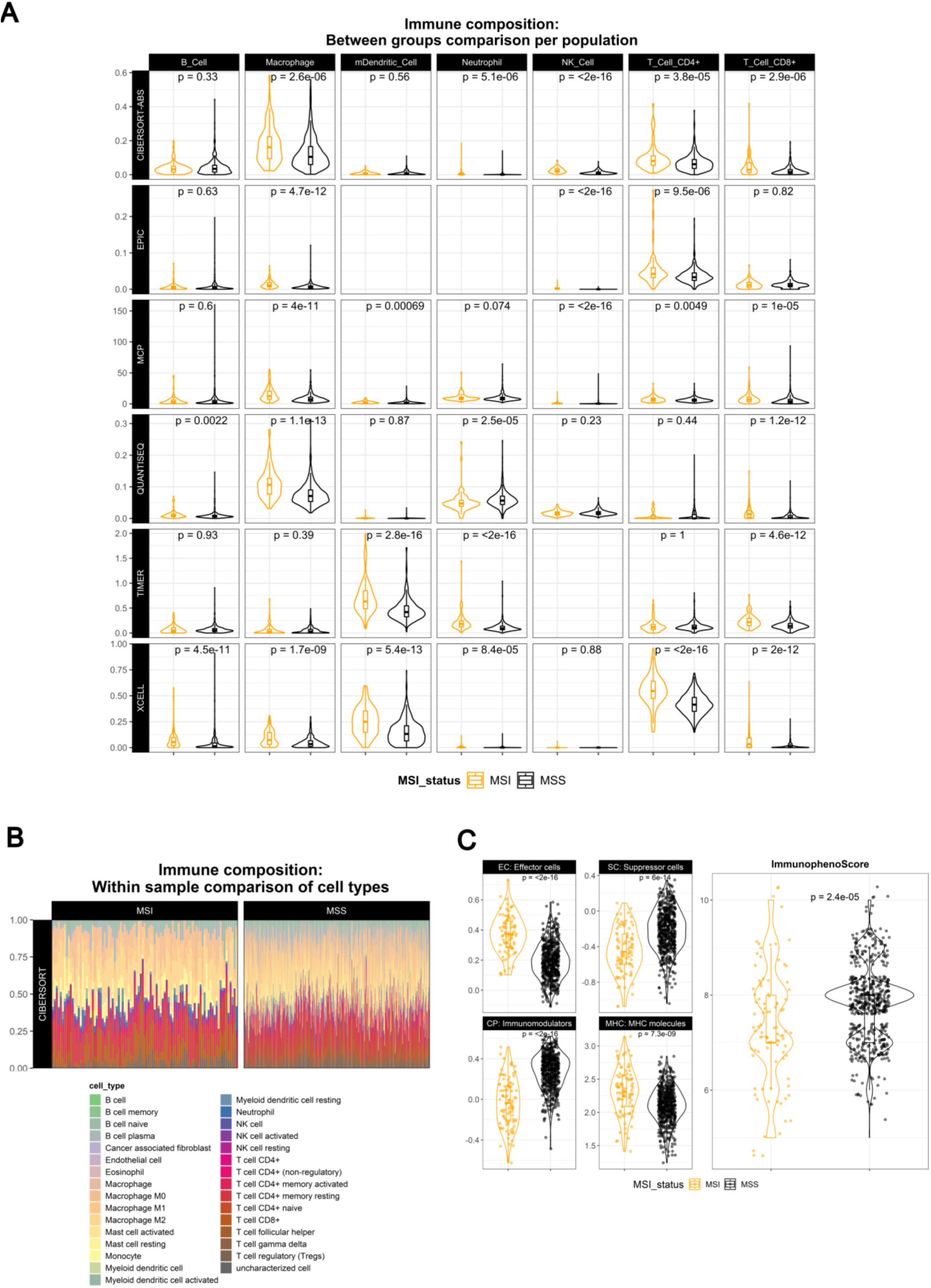
Immune composition of MSI and MSS colorectal tumor’s microenvironment. Immune deconvolution analysis from the Immune Composition (IC) module resulted in (**a**) boxplots ring MSI / MSS tumors for each major category generated by integrating the results from the odeconvR package; (**b**) a stacked barplot obtained by CIBERSORT showing the absolute ctions of each immune cell population within each sample, split by the MSI / MSS variable; plots obtained by the Immunophenoscore approach, comparing MSI / MSS tumors for the mmunophenoscore as well as for the four major determinants of immune response (Effector Suppressor Cells, Immunomodulators and MHC molecules).

## Discussion

In recent years, scientific research has witnessed a rapid evolution of high throughput technologies, which in turn led to a shift in focus towards data analysis over experimental procedures. High-throughput technologies and workflows have become an integral component of biological research, leading to considerable expansion of the bioinformatics field. The increasing availability of multi-omics datasets opens new opportunities for complex integrative data analysis, providing for a more thorough assessment of complex biological phenomena. However, the analysis of such large datasets can prove challenging, and it is becoming crucial to ensure that the ability to examine this type of data is not limited to individuals with advanced computer skills. With sequencing technologies becoming more ubiquitous in the experimental labs and clinical settings, it is vital to provide access to both easy-to-use and comprehensive open-source tools to overcome the barrier of ‘big data’ analysis and enable wet-lab as well as clinical scientists to analyze common omics data derived from their own experiments.

In this regard, GEGVIC provides a comprehensive R toolbox specifically designed to facilitate diverse bulk genomic and transcriptomic analyses in cancer research, catering to researchers with limited computational expertise. Its modular pipeline combines a variety of widely used methods, making it a versatile tool to analyze tumor mutation profiles, assess phenotypic- associated transcriptomic particularities, and infer the immunologic composition of the TME. To increase accessibility, we also developed a user-friendly web-based interface, GEGVICshine, which completely removes the coding barrier, rendering the tool accessible to scientists with no computer-programming skills. However, the flexibility remains for researchers with minimal R expertise to incorporate GEGVIC functions into their pipelines to expedite routine RNAseq and genomics data analyses. Also, the local R package version provides the advantage of not requiring a server to run analyses, thus eliminating computational constraints that may arise when working with large cohorts like the TCGA-COADREAD (n=632 samples from RNAseq, n=484 from WES).

GEGVIC offers various advantages over the current methods aimed at simplifying bioinformatics analysis. First, it allows for simultaneous analysis of genomic and transcriptomic data, making it suitable for investigating biological questions of different nature. Additionally, it provides greater flexibility in terms of input restrictions, statistical parameters and output customization. Moreover, GEGVIC automatically adapts its workflow to analyze mouse data with different versions of genomic annotations, without requiring additional interventions. The package also includes gene information from the Biomart database (12), which eliminates the need for users to convert gene identifiers to gene symbols as required in some steps of the pipeline. In terms of the scope of the proposed analyses, it is important to note that GEGVIC offers complementary analytical approaches at each module, allowing users to adopt different methodologies and establish a more solid basis to obtain meaningful results. For example, to perform pathway enrichment analysis, both GSEA (14) and GSVA (15) are proposed, facilitating the comparative analysis from different statistical perspectives (46,47). GSEA is the most commonly used method to compare a set of differentially expressed genes (supervised approach), but GSVA offers an alternative strategy that involves characterizing the degree of expression enrichment of gene sets in each sample, enabling an unsupervised comparison of gene sets activity between different samples from the groups of interest. In the context of immunological dissection of the TME, while both the *immunodeconv* and *immunophenoscore* strategies rely on assessing the expression levels of markers associated with distinct immune cells, utilizing multiple methods may enable to obtain more accurate and reproducible results, which can strengthen the interpretation and confidence of the analysis. Additionally, the IPS provides information not only about the anti-tumor immune infiltrates but also about the presence of immune suppressive cells (Tregs and MDSCs), which are indicative of tumor immune evasion (48,49). Finally, the *Genomic Variation* module offers, on the one hand, an overview of the mutational landscape identified in the studied dataset and comparison groups, including the identification of oncogenic ‘driver’ events, i.e., high frequency mutations with a key role in cancer development, and the calculation of the TMB, which is strongly associated with the amount of immunogenic neopeptides displayed on the tumor cell surface that influence patients response to immunotherapies (31,50). On the other hand, GEGVIC predicts mutational signatures for each sample, which are defined by the type of DNA damage and DNA repair processes that result in genomic alterations (33). Cancer genomes may carry several mutational signatures, representing the imprints of all the mutational processes that have occurred throughout cancer development. These signatures can provide the physiological readout of the biological history of a cancer and can distinguish ongoing mutational processes from historical ones (51).

One important limitation of GEGVIC is its lack of control over the quality parameters of the input. As a result, it is essential to subject the input files to initial quality control (QC) assessments before the analysis to prevent any potential artefacts or technical biases from influencing the final output. For expression analyses, quality issues on raw RNA-seq data can significantly distort analytical results and lead to erroneous conclusions (52,53). Quality monitoring of this data involves several steps from the raw reads, including the assessment of sequence quality, GC content and the detection of sequencing errors, PCR artifacts and contaminations. While some easy-to-use tools already exist for this purpose, such as RNA-QC-chain (54), their processing is computationally intensive and requires high performance computation not available in most experimental and clinical laboratories. Furthermore, the lack of clear consensus regarding the best analytical practices makes choosing the appropriate method for analysis a challenging task. We chose DESeq2 for DGE analysis over other popular options like edgeR (55) and limma-Voom (56), although no clear consensus exists regarding the optimal choice (52,57). In any case, due to the modular architecture of GEGVIC, incorporation of new features, methods and/or additional customization of the visualizations can be easily implemented to further improve and broaden the scope of the tool.

### Availability and requirements

Project name: GEGVIC

Project home page: https://github.com/oriolarques/GEGVIC, https://github.com/oriolarques/GEGVICshine, https://gegvic.vhio.net/

Operating system(s): Platform independent

Programming language: R and Shiny

Other requirements: R 2.10 or higher, but up to 4.1.3 or developmental version of R (4.3.0) License: MIT

Any restrictions to use by non-academics: None

### Availability of data and materials

All data and codes generated or analyzed during this study are included in this published article and its supplementary information files.

## List of Abbreviations

*CRC:*: Colorectal cancer
*DGE:*: Differential gene expression
*DNA-seq:*: DNA sequencing
*GE:*: Gene expression module
*GMT:*: Gene Matrix Transposed format
*GSEA:*: Gene set enrichment analysis
*GSVA:*: Gene set variation analysis
*GUI::*: Graphical user interface
*GV:*: Genomic variations module
*IC:*: Immune cell composition module
*IPG:*: Immunophenogram
*IPS::*: Immunophenoscore
*MSI:*: Microsatellite instability
*MSS:*: Microsatellite stability
*PCA:*: Principal component analysis
*RNA-seq:*: RNA sequencing
*TME:*: Tumor microenvironment
*VST:*: Variance stabilizing transformation

## Declarations

## Author’s contributions

O.A made the conception, data acquisition, coding, data analysis and writing the first draft. L.B made the conception, participated in data analysis and wrote the final version of the draft. All authors read and approved the final manuscript.

## Acknowledgments

We want to thank Dr. Héctor G. Palmer and Fundación Cellex for their institutional support, as well as to VHIO informatics support unit (AD Systems) for their contribution in maintaining the GEGVICshine server.

