## Supplementary Material for "GEGVIC: A workflow to analyze Gene Expression, Genetic Variations and Immune cell Composition of tumor samples using Next Generation Sequencing data"

#### Installation

The GEGVIC R package can be installed following the instructions from the GitHub page (https://github.com/oriolarques/GEGVIC). The Shiny app can be locally installed in R from the GitHub page (https://github.com/oriolarques/GEGVICshine), download the docker image (https://hub.docker.com/r/oarques/gegvicshine) or be used directly from the dedicated server https://gegvic.vhio.net/.

#### RNA-seq and mutational data

TCGA-COADREAD RNA-seq HTSeq raw counts were obtained from the NIH GDC Data portal (https://portal.gdc.cancer.gov/). Raw mutation annotation files were obtained from the FireBrowse repository (http://firebrowse.org/). Since the data was aligned to the hg19 version of the human genome, we used the liftOver tool from the UCSC Genome Browser (https://genome.ucsc.edu/cgi-bin/hgLiftOver) to convert the genomic coordinates to the hg38 version. Microsatellite stability status was also obtained from Firebrowse as part of the clinical data.

**Input format**

Gene expression and genomic input data was formatted to follow the requirements of the GEGVIC package. This are listed as follows:

- RNA-sequencing counts (expression matrix): Table containing gene counts as rows and samples as columns. The first column must contain gene identifiers that can be either *NCBI ID*, *ENSEMBL gene ID* or *HGNC ID* and its column name must be adequately named as either: *entrezgene_id, ensembl_gene_id hgnc_symbol*.

- Genomic variation data table (MAF file): Table containing short variant calls. Necessary columns must have the following names (following the MAF format, https://docs.gdc.cancer.gov/Data/File_Formats/MAF_Format/) *Hugo_Symbol* (Gene symbol from HGNC); *Chromosome* (Affected chromosome); *Start_Position* (Mutation start coordinate); *End_Position* (Mutation end coordinate); *Reference_Allele* (The plus strand reference allele at this position. Includes the deleted sequence for a deletion or "-" for an insertion); *Tumor_Seq_Allele2* (Tumor sequencing discovery allele); *Variant_Classification* (Translational effect of variant allele. Can be one of the following: Frame_Shift_Del, Frame_Shift_Ins, In_Frame_Del, In_Frame_Ins, Missense_Mutation, Nonsense_Mutation, Silent, Splice_Site, Translation_Start_Site, Nonstop_Mutation, RNA, Targeted_Region); *Variant_Type* (Type of mutation. Can be: 'SNP' (Single nucleotide polymorphism), 'DNP' (Double nucleotide polymorphism), 'INS' (Insertion), 'DEL' (Deletion)); *Tumor_Sample_Barcode* (Sample name).

- Samples metadata: A table containing additional information about the samples that can be used to create groups, such as '*Response*' or '*No-Response*' to a therapy. The first column must be named *Samples* and use the same sample nomenclature as in the RNA-sequencing matrix and genomic variation data tables.

**Differential gene expression (DGE) analysis**

To perform DGE analysis, GEGVIC uses functions from DESeq2 (1) to first read the raw count matrix and model design and then perform the appropriate calculations. The user can define the shrinkage method between the different options: *apeglm*, *ashr*, *normal,* or *none* to skip this step. The Principal Component Analysis (PCA) generated by this module is produced using the dedicated function from the package after normalizing and transforming counts using the variance stabilizing transformation (VST).

**Gene Annotation**

Association between Ensembl and/or NCBI gene IDs with the official gene symbols (HGNC: HUGO Gene Nomenclature Committee) is a feature offered by GEGVIC for the two more recent versions of the Homo sapiens (GRCh37 and GRCh38) and Mus musculus (GRCm38 and GRCm39) genomes. Annotation files were created using the biomaRt package (2). They contain the following information for each of the genomes: ensembl_gene_id, hgnc_symbol, entrezgene_id, transcript_length, refseq_mrna.

**Pathway Enrichment analysis**

Gene Set Enrichment Analysis (GSEA) is performed using the functions from the clusterProfileR v4.0 package (3), which include first the reading of the gene sets provided by the user in form of a gmt file and then GSEA computation. Downstream analyses are performed using the GSEAmining package (4) with default parameters.

Gene sets collection from MSigDB can be downloaded from the database page [http://www.gsea-msigdb.org/gsea/index.jsp](http://www.gsea-msigdb.org/gsea/index.jsp). Users can also create their own custom gene sets. Instructions on how to create a GMT file can be found in the database wiki page ([https://software.broadinstitute.org/cancer/software/gsea/wiki/index.php/Data_formats](https://software.broadinstitute.org/cancer/software/gsea/wiki/index.php/Data_formats)).

In order to perform Gene Set Variation Analysis (GSVA), the GSVA package is used (5), which allows the option of using different methods (GSVA, ssGSEA or Z-score). By default, the program uses the Hallmark collection of gene sets from the MSigDB (6), but the user can provide other gene sets, as commented before for GSEA, in the form of a gmt file. In this case, the GSEABase package (7) is used to read the file. Results are shown in a form of a clustered heatmap using the pheatmap R package (8).

**Immune cell prediction**

Prediction of immune cellular composition is achieved using the different algorithms (quanTIseq, TIMER, MCP-Counter, xCell, EPIC and CIBERSORT) that are executed using the immunedeconv R package (9). All methods are executed using default parameters. To use CIBERSORT the user needs to register on the CIBERSORT web page (https://cibersort.stanford.edu), obtain a license and download the source code in form of two files '*CIBERSORT.R*' and '*LM22.txt*'. Then, it is necessary to specify the path to the storage location of such files in the 'cibersort' argument. In any case, the Immune cell Composition module can be used without including CIBERSORT. Integration of different immune cells identified by each algorithm into the seven categories for the ic_plot_comp_samples() function was done grouping different subpopulations into the following groups: B-Cells, Macrophages (M0,M1 or M2), mDendritic_Cell (resting, activated or unspecified), Neutrophil, NK-Cell (resting, activated or unspecified), T-Cell_CD4+ (central memory, effector memory, activated, resting, or unspecified) and T-Cell_CD8+ (resting, activated or unspecified). Additional cell types were classified as Others (including Mast cells, Cancer associated fibroblasts, Endothelial cells and stem or progenitor cells). The complete table can be found in the github page of GEGVIC (https://github.com/oriolarques/GEGVIC/blob/main/data/ic_grouping.rda). To calculate immunophenoscore (IPS) and immunophenograms (IPG) we adapted the code from Charoentong et al. (10).

### Mutation evaluation

A summary of mutational data and an oncolplot are created by using the functionalities of the maftools R package (11). Mutational load is calculated as the total number of mutations per sample and the graphical representation is done using the ggplot2 R package. Mutational signatures are predicted using the deconstrucSigs R package (12). GEGVIC contains versions 2 and 3.2 of the different mutational signature matrices for single and double base substitutions for the different versions of the genome specified before. Those matrices were obtained from COSMIC (https://cancer.sanger.ac.uk/signatures/sbs/). Resulting plots are generated using ggplot2.

### Package versions

R (4.1.3), DESeq2 (1.34.0), GSEABase (1.56.0), GSEAmining (1.4.0), GSVA (1.42.0), SummarizedExperiment (1.24.0), apeglm (1.16.0), clusterProfiler (4.2.2), deconstructSigs (1.9.0), dplyr (1.1.0), ggplot2 (3.4.1), ggplotify (0.1.0), ggpubr (06.0), ggrepel (0.9.3), grid (4.2.0), gridExtra (2.3), immunedeconv (2.1.0), patchwork (1.1.2), pheatmap (1.0.12), rlang (1.0.6), tibble (3.2.0), tidyr (1.3.0), maftools (2.10.05).
