## Supplementary Figures for "GEGVIC: A workflow to analyze Gene Expression, Genetic Variations and Immune cell Composition of tumor samples using Next Generation Sequencing data"

**Supplementary Figure 1. Gene Expression Module.** ***Results obtained from a random subset of TCGA COAD-READ dataset (demo input).*** This module includes the (0) *Data Exploration* analysis of transcriptomic data through a PCA; the (A) visualization via Volcano Plot of the most significant genes obtained from the *Differential Gene Expression* (DGE) analysis; and the pathway enrichment analysis following two methodologies: the supervised *Gene Set Enrichment Analyses (GSEA)*, which incorporates different possibilities for data visualization (B1), and the unsupervised *Gene Set Variation Analysis (GSVA),* which plots a heatmap of normalized GSVA enrichment scores for the specified gene sets collection within the analyzed samples.

**Supplementary Figure 2. Immune Composition Module.** ***Results obtained from a random subset of TCGA COAD-READ dataset (demo input).*** This module includes the (A) *Immune Cell Composition* analysis of transcriptomic data obtained from the *ImmunodeconvR* package. Boxplots display the distribution of the scores obtained for each comparison group and method for each main immune category defined by GEGVIC, while the stacked barplots that can be obtained from CIBERSORT, EPIC and quanTIseq showing the absolute cell fractions of each immune cell population within each sample; and (B) *Immunophenogram / Immunophenoscore* analysis, which generates boxplots to compare the distribution of the scores obtained for each comparison group and category.

**Supplementary Figure 3. Genomic Variation Module.** ***Results obtained from a random subset of TCGA COAD-READ dataset (demo input).*** This module includes the (A) *Mutational landscape* analysis, which generates a comprehensive overview of the most common genomic alterations identified in the cohort as well as the mutational load calculated for each sample; and (B) the *Mutational Signature* analysis that calculates the weight of each mutational signature in each sample, which can be visualized by the stacked barplot or by a heatmap.

### Gene Expression Module

#### 0. Data exploration

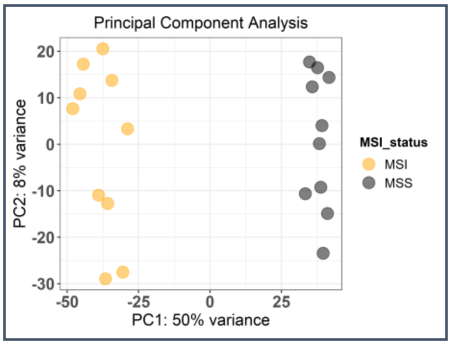


#### A. Differential Gene Expression (DGE) analysis

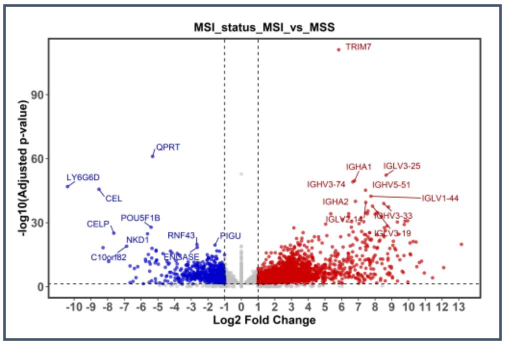


#### B1. Gene Set Enrichment Analysis (GSEA)

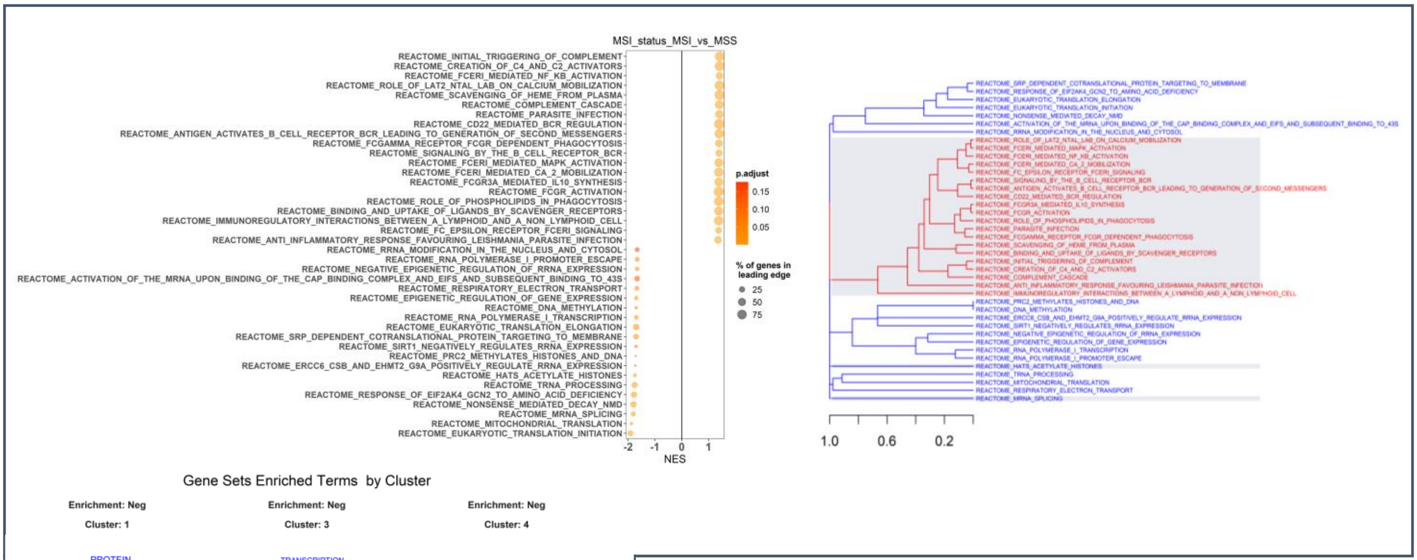


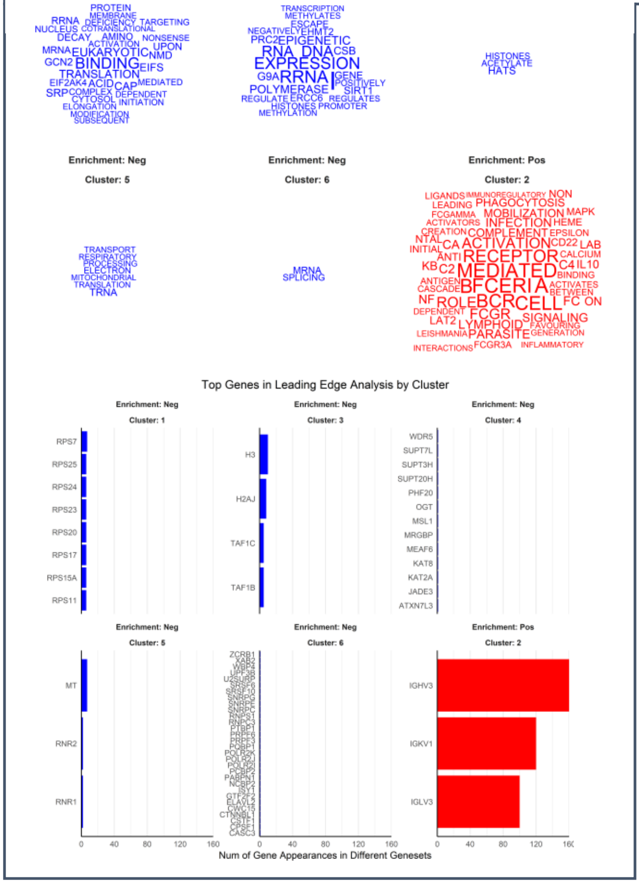


#### B2. Gene Set Variation Analysis (GSVA)

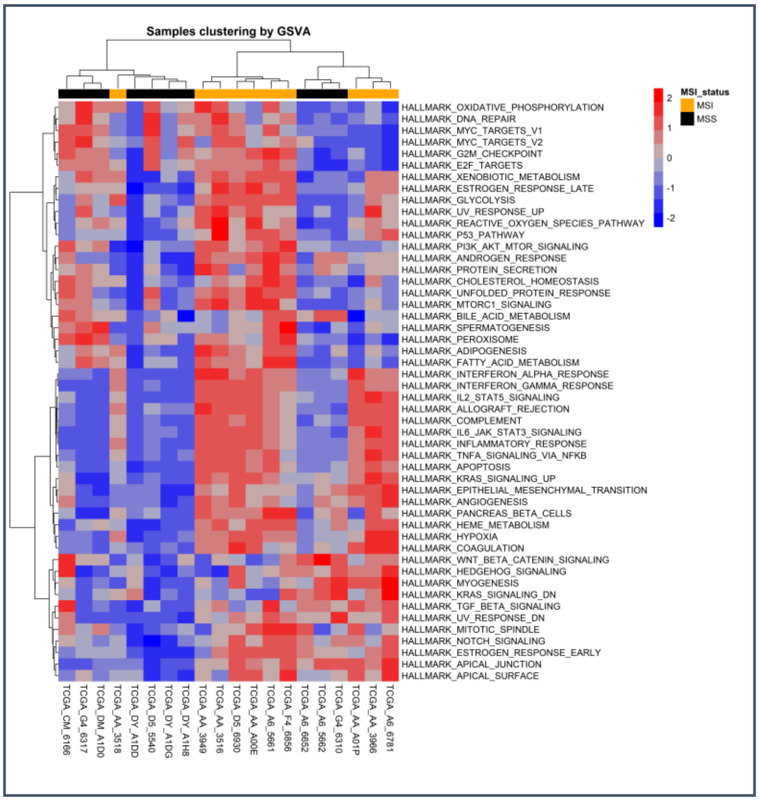

### Immune Composition Module

#### A. Immune Cell Composition

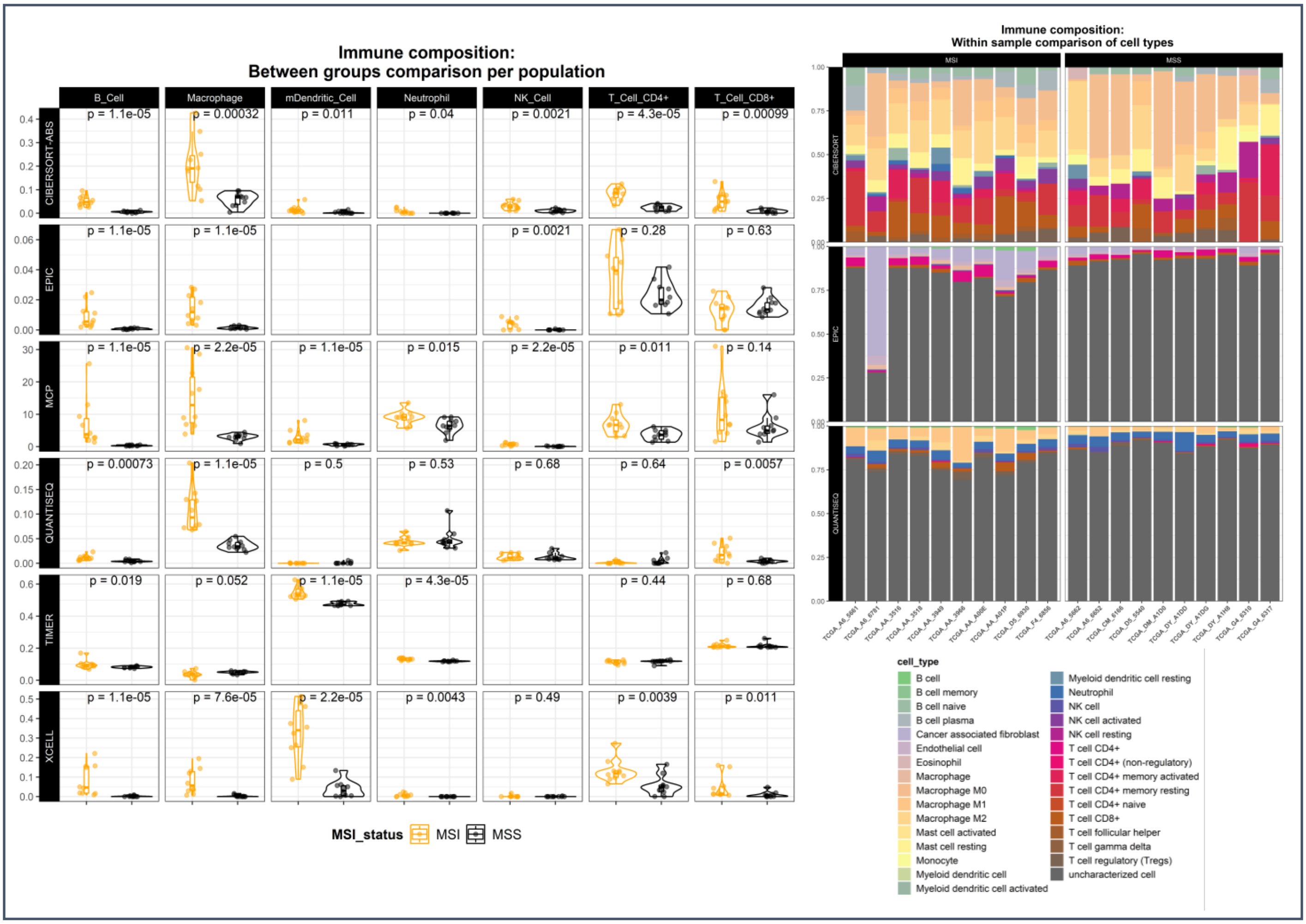


#### B. Immunophenogram / Immunophenoscore

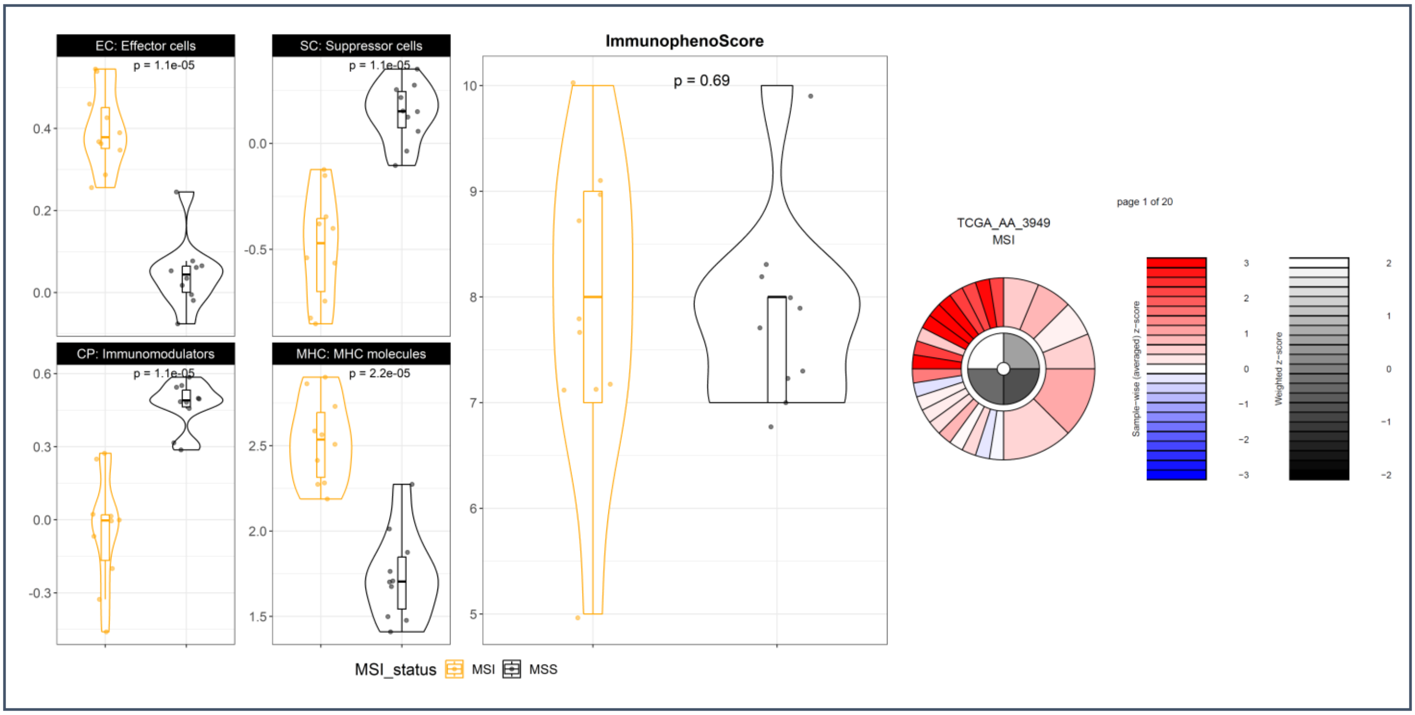

### Genomic Variation Module

#### A. Mutational Landscape

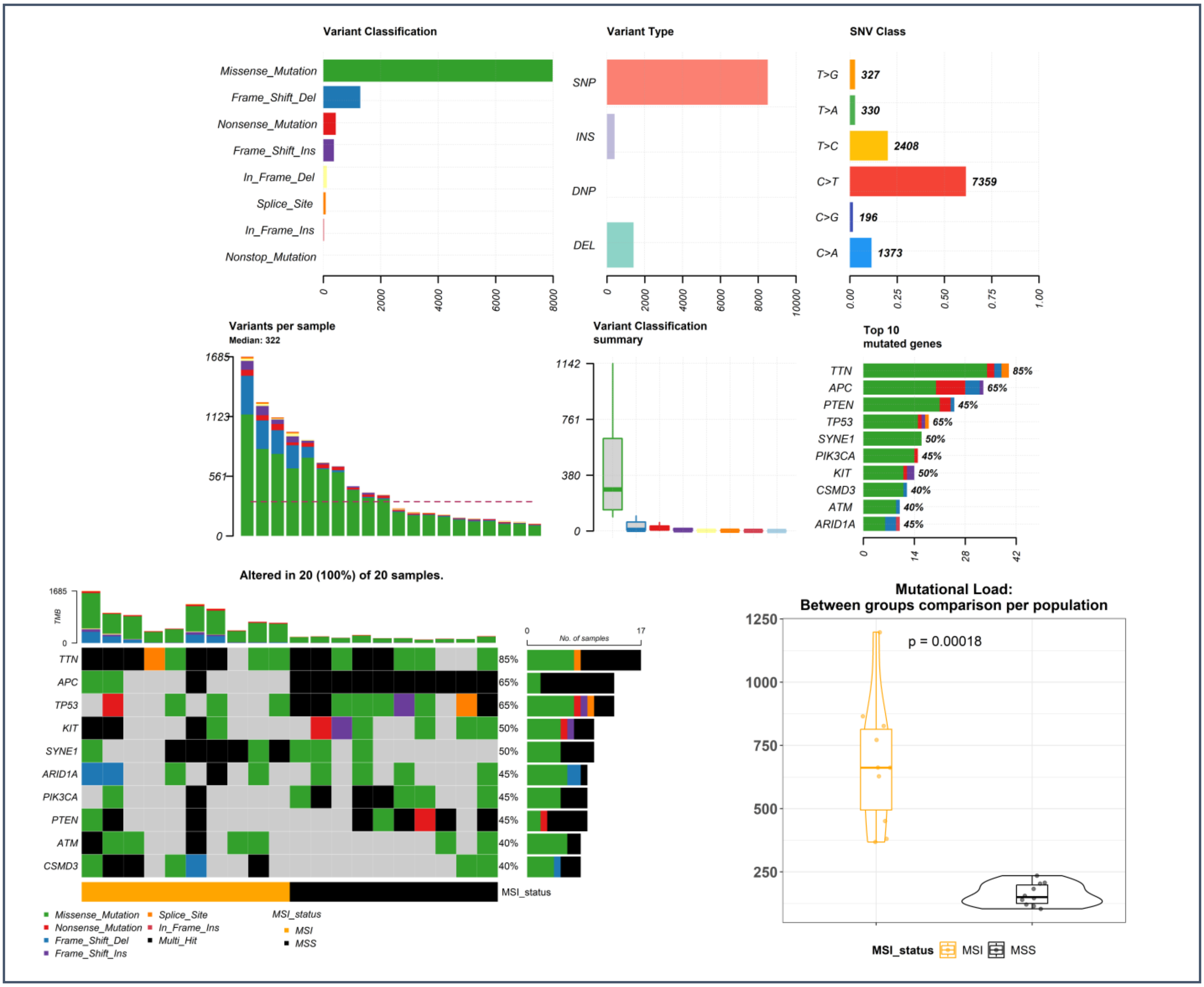


#### B. Mutational Signatures

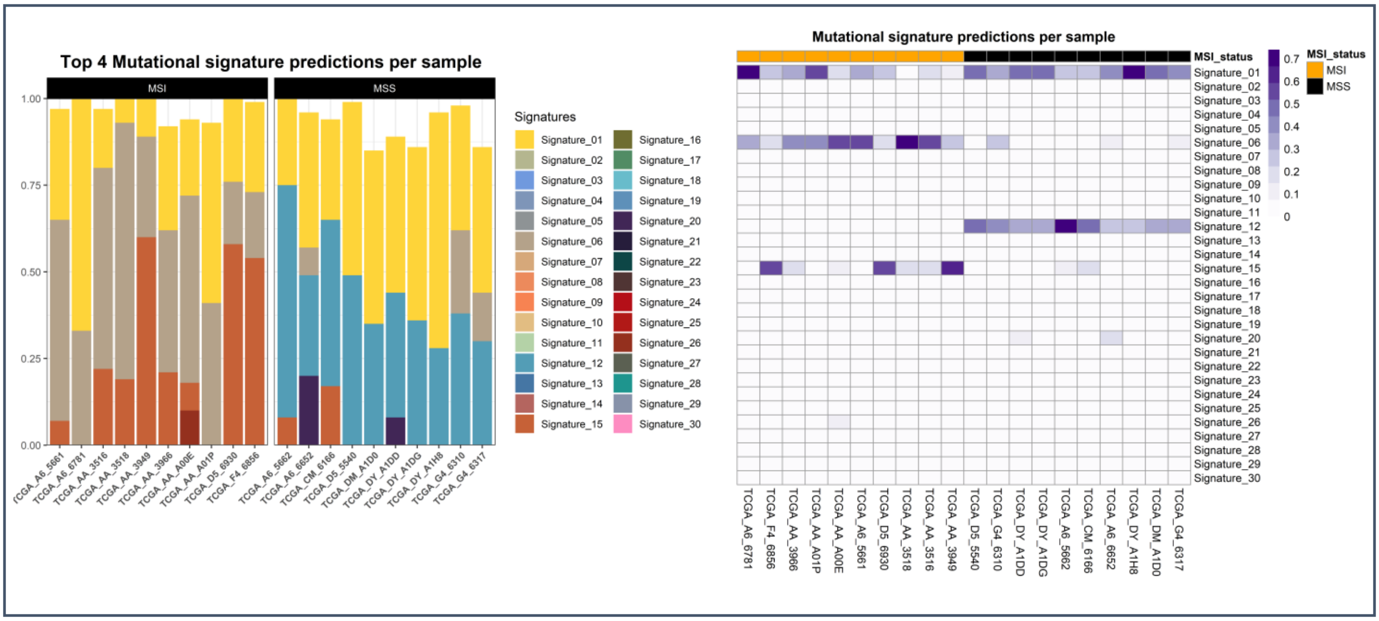
